# Smelling the Risk: Early Olfactory Deficits, Brain Networks, and Blood Markers of Alzheimer’s Disease Risk in Humanized APOE Mice

**DOI:** 10.1101/2025.06.06.658383

**Authors:** Hae Sol Moon, Zay Yar Han, Robert J Anderson, Ali Mahzarnia, Jacques Stout, Andrei Niculescu, Jessica Tremblay, Alexandra Badea

**Author notes:** <u>Contact</u> Alexandra Badea, PhD Quantitative Imaging and Analysis Labs. 161G Bryan Research Building, 311 Research Dr. Duke University Medical Center Durham, NC 27710. **Funding** We are grateful for NIH support through RF1 AG057895, R01 AG066184, RF1 AG070149.

## Abstract

Olfactory impairment is a hallmark of early Alzheimer’s disease (AD), but the underlying mechanisms connecting sensory decline to genetic and environmental risk factors remain unclear. Our integrative analysis combines ethologically relevant olfactory behavior assays, high-resolution diffusion MRI connectomics, and blood transcriptomics in a large cohort of humanized APOE mice stratified by APOE genotype (APOE2, APOE3, APOE4), age, sex, high-fat diet, and immune background (HN).

Behaviorally, APOE4 mice exhibited accelerated deficits in odor salience, novelty detection, and memory, especially when exposed to a high-fat diet, whereas APOE2 mice showed resilience (ANOVA: APOE x HN, F(2,1669)=77.25, p<0.001, eta squared effect size = 0.08). Notably, age and diet exerted compounding effects, with older and HFD-fed mice displaying reduced odor-guided exploration (diet x age: F(1,1669)=16.04, p<0.001, eta squared effect size = 0.01). Memory analyses revealed robust genotype- and age-dependent impairments: at 24- and 48-hour delays, recognition indices were significantly lower in APOE4 mice compared to APOE2 (long-term memory: APOE x HN, F(2,395)=5.6, p=0.004).

Elastic Net-regularized multi-set canonical correlation analysis (MCCA) linked behavior to brain network substrates, revealing subnetworks whose connectivity explained up to 24 percent of behavioral variance (sum of canonical correlations: 1.27, 95% CI [1.18, 1.85], p<0.0001). High-weighted connections between the ventral orbital cortex, somatosensory cortex, and cerebellar-brainstem pathways were identified as critical nodes for risk or compensation.

Integrative blood transcriptomics revealed eigengene modules strongly correlated with imaging changes in olfactory-memory circuits (for example, eigengene 2 vs. subiculum diffusivity: r = -0.5, p < 1e-30, explaining up to 24 percent of variance). Gene ontology analysis pinpointed shared pathways in synaptic signaling, translation, and metabolic regulation across brain and blood. Notably, glutamatergic and synaptic pathways were enriched among genes linking peripheral and central compartments.

Collectively, these results demonstrate that olfactory behavior, quantitatively shaped by genotype, age, diet, and immune status, serves as a sensitive and translatable early biomarker of Alzheimer’s disease risk. Our systems-level approach identifies specific brain networks and peripheral molecular signatures underlying sensory-cognitive vulnerability, providing a robust framework for early detection and targeted intervention in AD.

## Introduction

Alzheimer’s disease (AD) is a complex and multifactorial neurodegenerative disorder that primarily affects the aging population and remains irreversible, with no definitive cure to date [1]. It is marked by a progressive deterioration of cognitive functions, including memory deficits and behavioral changes, which mirror the accumulation of neuropathological alterations across the brain. Hallmark features of AD pathology include the aggregation of amyloid-β plaques, hyperphosphorylated tau tangles, cortical and subcortical atrophy, and the disruption of neural circuits [2–4]. The disease unfolds gradually over many years with a prolonged preclinical phase during which subtle physiological and behavioral abnormalities emerge about 10 to 15 years before obvious symptoms and possible clinical diagnosis [5]. This extended prodromal window underscores the critical need for sensitive and reliable early biomarkers, predictive of cognitive decline, and capable of identifying individuals at risk. Determining the initial circuit-level perturbations that may precede irreversible neurodegeneration [5, 6] may offer new targets for more effective interventions.

One promising target is the olfactory circuit, as olfactory dysfunction has been proposed as a low-cost, and accessible biomarker for early-stage detection [7, 8]. Individuals at risk for or in the early stages of AD often demonstrate impairments in odor detection, discrimination, identification, and memory well before the onset of clinically diagnosable cognitive decline [9, 10]. This early vulnerability is partly attributable to the unique anatomy of the olfactory system, which is directly and densely connected to key brain regions implicated in AD pathology including the entorhinal cortex, amygdala, and hippocampus [11]. These regions are among the first to undergo neurodegenerative changes and pathological protein accumulation in both human patients and animal models of AD [11, 12]. In clinical research, olfactory function is commonly assessed through standardized cognitive tests such as the University of Pennsylvania Smell Identification Test (UPSIT), Sniffin’ Sticks, and AROMHA Brain Health Test (ABHT) [8, 13, 14]. These tasks not only measure perceptual sensitivity but also engage higher-order memory systems, making them particularly informative for probing early sensory-cognitive decline [15]. Given the close functional and anatomical relationship between olfaction and memory, olfactory testing provides a uniquely sensitive lens through which to investigate the earliest neural disruptions associated with AD [16]. However, while olfactory behavior is known to involve networks of brain regions, the relationship between behavior performance on these odor based tasks and brain structural connectivity remains poorly understood in the context of AD, and AD risk. A comprehensive framework that integrates relevant olfactory tasks with systems-level neuroimaging—such as diffusion MRI based connectomics—could uncover new insights into the early sensory-cognitive changes in AD and their neural circuits that underlie them.

The clinical relevance of olfactory dysfunction in AD has spurred the refinement of mouse behavioral assays, yet key methodological hurdles remain. For example odor memory is long-lasting but can easily be masked by dominant visual cues or by odors that are insufficiently novel [17]. Still, animal models can help advance our understanding of AD, particularly in dissecting the complex interplay between genetic and environmental risk factors[18], and their impact on behavior and associated brain circuits. Mouse models, in particular, provide a controlled experimental platform that allows for precise manipulation of variables that are difficult to isolate in human populations like diet [18–21], balancing population based the carriage of different APOE alleles, age, or diet.

This level of experimental control enables causal inferences and the ability to investigate mechanistic underpinnings of disease-related phenotypes. Mice expressing the human APOE4 allele, model the highest known genetic risk for AD, relative to the control APOE3 allele, and the protective APOE2 alleles [22]. APOE4 mice has been found to have olfactory memory impairments compared to APOE4 allele mice [23]. Carrying the APOE4 allele is known to affect olfactory-related brain regions, which are intricately connected to areas involved in motivated behavior and cognition through shared neural circuits [24]. Moreover, it is possible to get detailed brain tractography in mice by using high-resolution ex-vivo MR imaging to study subtle connectivity changes related to brain function and structure that may precede overt neurodegeneration in aging and AD pathology[25, 26]. These features make preclinical models uniquely suited for testing hypotheses about the earliest contributors to AD pathophysiology and for evaluating biomarkers changes under tightly regulated conditions.

There have not been many studies on how the decline in brain regions related to olfaction in normal individuals predicts later disease, and how these regions are connected to other regions. This motivates our study to find whether brain connections related to olfaction change in image related properties, and how these changes relate to olfactive behavioral performance especially in long-term memory [27], and whether these changes can be captured by peripheral markers.

We aim to investigate how AD-related genetic and environmental, modifiable risk factors impact olfactory behaviors in mice, and how these behavioral phenotypes relate to alterations in brain connectivity. Specifically, we test odor preference, reaction to novel odors, and memory—in mice that model various combinations of AD risk factors. These mouse models include humanized variants of the apolipoprotein E (APOE) gene (ε2, ε3, and ε4), the strongest genetic risk factor for late-onset AD in humans; humanized nitric oxide synthase 2 (hNOS2) to mimic human-like immune responses; sex (male and females); age (young and middle-aged); and environmental stressors such as high-fat diet environment [28–30]. By studying these factors both independently and in combination, we aim to identify not only which individual risk variables impair olfactory performance, but also which synergistic combinations lead to more pronounced olfactory behavioral decline. To connect behavior with brain circuitry, we pair this behavioral assessment with high-resolution ex vivo diffusion MRI-based connectomics[31]. This allows us to map the structural architecture of the brain and quantify how connectivity patterns relate to performance on odor-related tasks. We seek to identify specific brain regions and networks which structural integrity correlates with olfactory behavior, especially in circuits implicated in memory and sensory integration. Ultimately, our goal is: (1) to determine how genetic and environmental risk factors for AD impact odor-related behavioral markers, (2) to uncover the brain circuits that underlie these behavioral changes, thereby offering insights into the early pathophysiology of Alzheimer’s disease, and (3) to investigate mechanistic changes in peripheral markers that can offer a noninvasive insight into central processes.

Finally we assessed whether regional differences in brain gene expression can be mapped using peripheral blood transcriptomics. Our findings show that we can detect such central marker changes using peripheral markers, and this method offers rich insight into pathways altered in individuals with different risk profiles for AD. We hope our study will contribute to the development of translatable early biomarkers and add methods to improve our mechanistic understanding of how diverse risk factors interact to shape disease trajectories.

## Methods

### Animals

Animal procedures were approved by the Duke Institutional Animal Care and Use Committee. To model genetic risk for late-onset Alzheimer’s disease (LOAD), we used animals lacking the endogenous mouse Apoe gene and homozygous for one of the three major human APOE alleles: the high-risk APOE4, the neutral APOE3, and the protective APOE2. These lines were crossed with mice where the mouse Nos2 gene was replaced by the human NOS2 gene (HuNOS2tg/mNos2−/−, herein termed HN). This replacement models the reduced NOS2 expression seen in human macrophages compared to murine counterparts, aligning redox homeostasis more closely with the human. Both sexes were included. Mice were naturally aged, with dietary exposures to either regular chow (2001 Lab Diet) or a high-fat diet (HFD; D12451i, Research Diets) for the last 4 months of life. Control diet consisted of 13.6 kcal% fat, 57.5 kcal% carbohydrate (3.25 kcal% sucrose), and 28.9 kcal% protein, while HFD contained 45 kcal% fat (39 kcal% from lard; 5 kcal% from oil), 35 kcal% carbohydrate (17 kcal% sucrose), and 20 kcal% protein. Mice had ad libitum access to food and water.

The composition of the animal cohorts used for behavioral testing is detailed in **Table 1**. A total of 465 mice were included in the behavioral analysis, stratified by APOE genotype (APOE2, APOE3, APOE4), diet (control or high-fat diet), sex, age group (younger: ∼12 months, older: ∼18 months), and NOS2 genetic background (mouse NOS2 [mNOS2] or human NOS2 transgene [HN]). A subset of 173 mice was used for imaging, as described in [32]. RNA-seq was performed on tissue samples from up to 34 animals per brain region, specifically: hypothalamus (n=34), cingulate cortex (n=32), frontal cortex (n=32), hippocampus (n=27), amygdala (n=8), brainstem (n=7), caudate putamen (n=8), thalamus (n=8), olfactory bulb (n=7), substantia nigra (n=6), colliculus (n=6), and cerebellum (n=3), which were compared with blood samples from the same animals, to examine patterns of shared variation. The cohort was balanced across APOE genotype, sex, diet, age range, and NOS2 status.

**Table 1.** Behavioral Cohort Distribution by Genotype, Diet, Sex, Age Group, and NOS2 Background.

| APOE | Diet | Female | Male | 18 Months | 12 Months | mNOS2 | HN | Total |
| --- | --- | --- | --- | --- | --- | --- | --- | --- |
| **APOE2** | Control | 46 | 48 | 51 | 43 | 48 | 46 | 94 |
| **APOE2** | HFD | 37 | 34 | 31 | 40 | 29 | 42 | 71 |
| **APOE3** | Control | 45 | 41 | 41 | 45 | 43 | 43 | 86 |
| **APOE3** | HFD | 35 | 33 | 32 | 36 | 30 | 38 | 68 |
| **APOE4** | Control | 41 | 35 | 28 | 48 | 43 | 33 | 76 |
| **APOE4** | HFD | 39 | 31 | 23 | 47 | 34 | 36 | 70 |

## Behavior Assessments

### Odor Preference and Habituation–Dishabituation Test

All mice were acclimated to the testing room for 30 minutes prior to the start of behavioral testing. On Day 1, each mouse was placed in the center of a square Plexiglas open-field arena (40 × 40 cm) and allowed to explore freely for 10 minutes. Between subjects, the arena was wiped with 70% ethanol and water. We recorded and analyzed locomotor activity using (ANY-maze, Stoelting Co., Wood Dale, Illinois, USA) with overhead cameras. Following this, we assessed odor preference and olfactory sensitivity using a series of trials involving different dilutions of lemon scent in mineral oil, as well as mineral oil alone. The odor stimuli were presented within custom-designed, 3D-printed hollow objects featuring perforations to allow scent dispersion. Food-grade lemon essence was diluted in mineral oil at concentrations of 0 (vehicle), 0.001, 0.01, and 0.1. Each test lasted 2 minutes, during which four odorized objects were simultaneously presented. To reduce potential location bias and cross-contamination, the arenas were cleaned thoroughly between trials, and the positions of the objects were pseudorandomized between sessions. Each condition was tested in four consecutive trials, followed by a fifth dishabituation trial, where the familiar odor (lemon) was replaced with a novel odor (vanilla or almond, depending on cohort) to assess novelty detection. Preference ratios were recorded based on the time spent investigating each object, and dishabituation indices were calculated as shown later. Exploration time was scored using ANY-maze behavioral tracking software (Stoelting Co., Wood Dale, IL, USA), a validated system for automated behavioral tracking in rodents [71], with experimenters blinded to group identity. Arenas and objects were custom-designed in-house to allow modular scent delivery while controlling for shape and texture.

### Novel Odor Memory

We analyzed odor-based memory tests using a protocol that began with two identical odors, followed by the introduction of a novel scent after specific time delays. Mice were first exposed to two objects scented with coconut oil across three identical habituation trials. To assess short-term memory, one coconut-scented object was replaced in trial 4 with an almond-scented object after a 1 hour delay. For long-term memory (24 hour delay), the novel scent was changed to anise, and for remote memory (48 hour delay), banana was used, as illustrated in **Figure 1C**. Across all tests, the positions of the familiar and novel odor objects were alternated to reduce spatial bias and isolate olfactory-driven responses, following procedures adapted from [33, 34].

**Figure 1.**
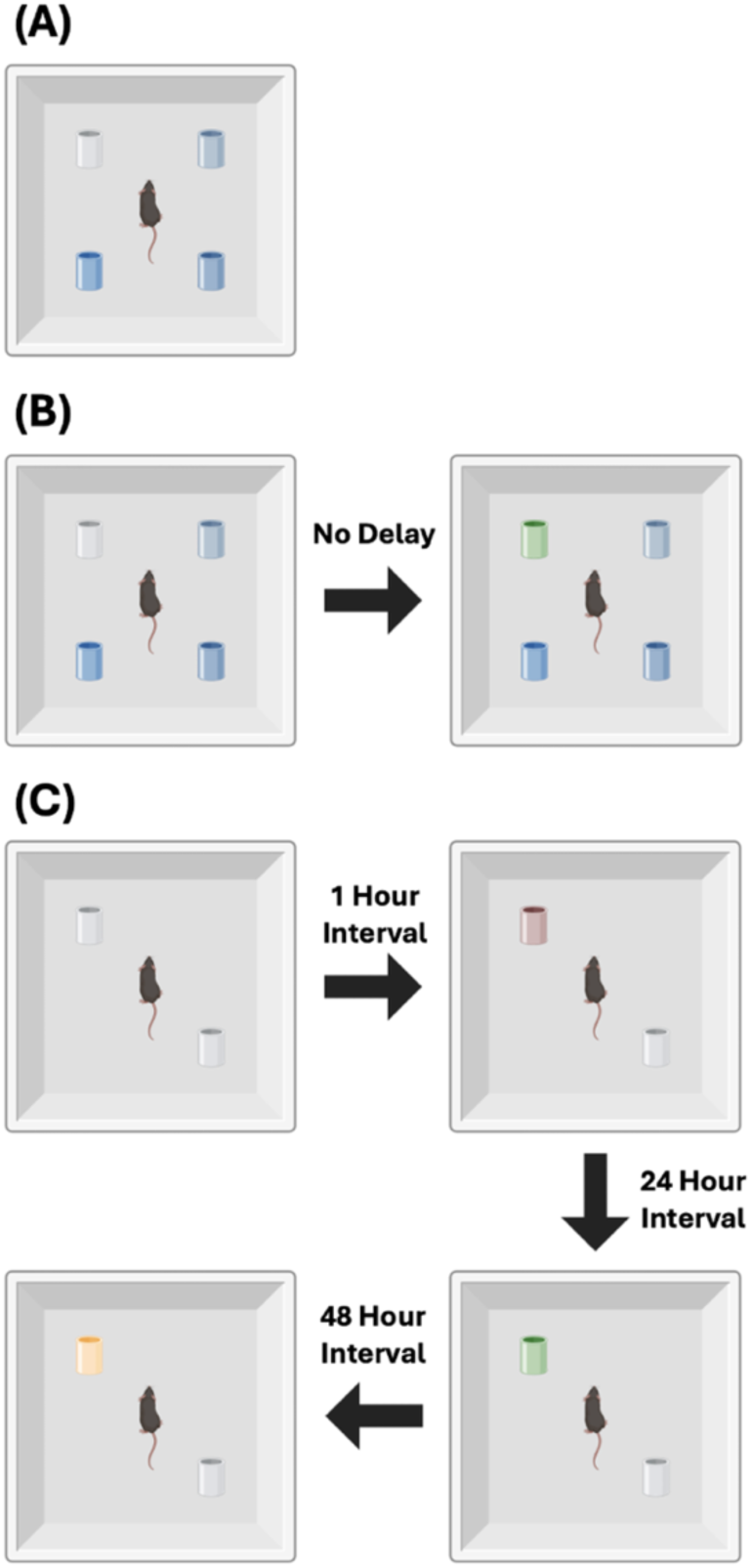
Olfactory Experiments Design. (A) *Odor preference during habituation:* Mice freely explored an open field containing four objects, each infused with different concentrations of lemon odorant. (B) *Habituation–Dishabituation task:* Mice were exposed to four consecutive trials with the same odor (lemon), followed by a dishabituation trial in which the lemon-scented object was replaced with one scented with vanilla. (C) *Memory tests:* To assess short-term, long-term, and remote olfactory memory, mice were first exposed to two objects containing the same novel odor. At 1 hour, 24 hours, and 48 hours after initial exposure, one of the odors was replaced with a new one to test recognition.

The following metrics were used to quantify behavior:

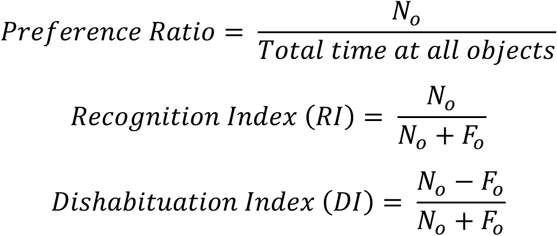

where *N_o_* is the investigation time at the novel odor, and *F_o_* is the time at the familiar odor.

### Statistical Analysis

Odor-exploration data were analyzed in R using either linear models (lm) or mixed-effects models (lme4 + lmerTest), with trial number or odor concentration as within-subject factors and APOE genotype, diet, immune background and sex as between-subject factors. Mouse identity was included as a random effect to account for repeated measures. For each memory stage, we first fit a linear model to assess the impact of individual traits on the recognition index; we then used a linear mixed-effects model—with memory type as an additional stage factor—to examine how these behavioral responses varied across memory durations and mouse traits.

Habituation was assessed by modeling a decline in exploration time across repeated trials. Dishabituation was evaluated by comparing exploration time in the final repeated odor trial, to the last trial presenting a novel odor, using a categorical predictor. Additionally, a dishabituation index was calculated and analyzed using standard linear models (lm), as each subject contributed a single index value. Post hoc comparisons were adjusted using Tukey’s HSD method via the emmeans package. Statistical significance was set at *p* < 0.05.

### Perfusion and Tissue Preparation for Imaging

After conclusion of the behavior tests, animals were sacrificed to prepare specimens for ex vivo imaging. Animals were euthanized under a surgical plane of anesthesia (100 mg/kg ketamine and 10 mg/kg xylazine) and perfused transcardially via the left ventricle with 0.9% saline (∼8 mL/min for 5 min), followed by 10% neutral buffered formalin phosphate containing 10% Gadoteridol (ProHance, Bracco Diagnostics) at the same flow rate and duration. Brains were left within the skull to preserve anatomical integrity and stored in formalin for 12 hours, followed by rehydration in 0.5% Gadoteridol in phosphate-buffered saline at 4°C. Heads were trimmed of extraneous tissue and placed in MRI-compatible tubes filled with perfluoropolyether (Galden Pro) for susceptibility matching.

### Image Acquisition and Processing

Diffusion-weighted MRI was performed on a 9.4T high-field scanner, as described in [32], using a 3D spin-echo sequence (TR/TE = 100 ms/14.2 ms, matrix = 420×256×256, FOV = 18.9×11.5×11.5 mm³, resolution = 45 μm isotropic). The protocol included 46 diffusion directions across two shells (b=2000 and 4000 s/mm²) and five b0 images. Acquisition was accelerated with 8-fold compressed sensing. Diffusion tensor metrics (e.g., fractional anisotropy), orientation distribution functions, and tractograms were reconstructed using MRtrix3[35], including streamline filtering via SIFT. Brain regions were segmented into 332 regions using the SAMBA pipeline[36], and whole-brain structural connectomes were constructed as adjacency matrices representing streamline counts between region pairs.

### Transcriptomics

RNA was extracted from blood and 12 select brain regions using QIAGEN kits. RNA-Seq experiments were conducted at the Duke Sequencing and Genomic Technologies Shared Resource Core to identify genes differentially expressed concerning age and sex. The RNA quality was assessed with a NanoDrop 2000 spectrophotometer. Samples were processed on an Illumina NovaSeq600 S2 platform, with 50 bp PE full flow cell, utilizing NuGEN mRNA-Seq with Any Deplete Globin. Sequencing data underwent quality control through FastQC, adhering to a Phred score cutoff above 20. As no adapter sequences were found adjacent to transcript reads, data trimming was not required. We aligned raw RNA-seq reads to a Mus musculus reference transcriptome, and the Salmon program [37] was utilized to quantify transcript abundance, as described in [38]. We performed GO enrichment on the top-loading genes for each principal component (PC) of our blood-based or brain based PCA in mouse. For each PC, genes were ranked by the absolute value of their loading and the top 200 retained. Mouse gene symbols were mapped to Entrez IDs via org.Mm.eg.db, and any PC with fewer than 10 mapped IDs was omitted. To identify biological processes associated with PC modules, we performed gene ontology (GO) enrichment analysis using the clusterProfiler package in R, querying the GO Biological Process (GO:BP) category. Multiple testing correction was applied using the Benjamini–Hochberg False Discovery Rate (FDR) method. All p-values reported for pathway enrichment are FDR-corrected. Pathways with an FDR-adjusted p-value < 0.05 were considered statistically significant. Only pathways passing the significance threshold were included in further analyses or visualization.

### Multivariate Modeling

To examine shared variance between brain structural connectivity and multidomain traits, we implemented a two-stage multivariate pipeline combining Elastic Net regression and Multi-set Canonical Correlation Analysis (MCCA), expanding upon previous work [38].

### Elastic Net Feature Selection

Dimensionality reduction was first performed using Elastic Net regularization [39]. The connectome matrix X (nodes × edges) was regressed onto behavioral outcomes Y using a penalty that blends L1 and L2 norms:

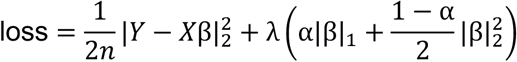

Here, β is the vector of coefficients to be learned, λ is the regularization parameter, and α (set to 0.5) balances sparsity (L1 norm) and ridge penalty (L2 norm). The optimal λ was identified using 10-fold cross-validation, retaining only those connectivity features with non-zero coefficients. This step ensured robust dimensionality reduction prior to multivariate modeling.

### Multi-set Canonical Correlation Analysis (MCCA)

We then applied MCCA to identify latent dimensions that link three domains: (1) structural connectomes, (2) olfactory-guided behavioral responses (odor salience, anhedonia, and odor memory), and (3) stratifying Alzheimer’s disease risk traits (APOE genotype, diet, immune background, age, and sex). MCCA finds maximally correlated linear projections across datasets by solving:

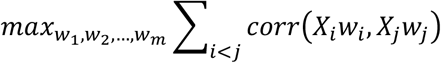

where X(k) is the standardized data matrix for domain k, and w(k) is its canonical vector. Each matrix was scaled and entered separately to preserve within-domain structure while allowing cross-domain alignment. The resulting canonical scores were back-projected into brain space by expanding feature indices to the original connectome matrix, enabling the identification of brain subnetworks that contribute most strongly to cross-domain variance.

This integrative approach allowed us to extract distributed network-level signatures of AD risk and behavior, reflecting early disruption of brain–olfactory-guided behavior alignment across systems.

### Peripheral Molecular Mechanisms Associated with Brain Imaging Features

To gain insight into molecular mechanisms underlying image feature changes we integrated complementary data streams in the same cohort of animals: 1) regional diffusion-MRI and volumetric metrics extracted from 3D structural images, and 2) blood-derived gene-expression modules. MRI acquisitions included high-resolution diffusion-weighted scans from which we computed standard tract-based metrics (fractional anisotropy, axial/radial/mean diffusivity) as well as the MAP-MRI indices (mean squared displacement, non-Gaussianity, q-space inverse variance, return-to-origin probability). Volumes were calculated as percent of intracranial volume, following brain segmentation and parcellating each brain into 332 anatomically defined regions, using SAMBA [36]. For focused analysis of olfactory-memory circuits, we selected eight a priori regions: the Piriform Cortex, Amygdalopiriform Transition Area, Posterolateral Cortical Amygdaloid Area, Perirhinal Cortex, Hippocampus, Postsubiculum, Parasubiculum, and Ventral Orbital Cortex.

Whole-blood transcriptomes were normalized before deriving 10 principal components—termed the eigengene—as summaries of expression measures. To assess gene–imaging coupling, we merged imaging and transcriptomic data with one row per subject–region–metric–eigengene combination. Within each brain region, both imaging metrics and eigengenes were z-scored, allowing regression slopes to be interpreted as Pearson correlation coefficients. Linear models were fit for each of the 560 metric–region–eigengene combinations (8 regions × 10 eigengenes × 7 imaging metrics), modeling standardized eigengene expression as a function of standardized imaging metrics. P-values were corrected using the Benjamini–Hochberg procedure to control the false discovery rate (FDR), and only associations with FDR < 0.05 were retained for downstream analysis.

For visualization, we selected the top five regions per metric–eigengene pair with the strongest correlations. We plotted signed effect sizes (β), –log_10_(FDR) values, and absolute correlations across a grid of metric × eigengene facets. All analyses were conducted in R 4.2 using the tidyverse, broom, and purrr packages. For functional interpretation, we performed Gene Ontology (GO) Biological Process enrichment using the top 100 genes from each module with the gprofiler2 package (organism: Homo sapiens), without initial p-value filtering to capture both strong and moderate associations. Results were processed to flatten list columns and visualized using ggplot2, displaying the top 10 enriched pathways per module based on –log_10_(p-value). All enrichment outputs were organized in a dedicated results directory for interpretation and figure generation.

To identify systemic molecular signatures associated with regional brain vulnerability in Alzheimer’s disease (AD), we analyzed the correlation structure between gene expression principal components (PCs) from blood and brain regions (hypothalamus, hippocampus, frontal cortex, and cingulate cortex). For each region, the top PCs from the brain dataset were correlated against the top PCs from the blood dataset using both Pearson and Spearman correlation tests. All raw p-values from these pairwise tests were adjusted for multiple comparisons using the Benjamini–Hochberg procedure. Brain–blood PC pairs with raw Pearson p < 0.05 were considered significant. For each significant brain–blood PC pair, we selected the top 200 genes with the highest absolute loading values in each PC. The intersection of these two gene sets (brain and blood) was used for Gene Ontology (GO, Biological Process) enrichment analysis, performed using the clusterProfiler package (mouse: org.Mm.eg.db; human: org.Hs.eg.db; keyType = “SYMBOL”; BH adjustment; p < 0.05). Only intersections comprising at least five genes were considered for enrichment testing. Enriched GO terms and their constituent genes were recorded for downstream interpretation. All computational steps, including correlations, FDR correction, and GO enrichment, were parallelized to expedite processing across regions.

This intersection-based approach allows for the identification of molecular pathways that are systemically reflected across both blood and brain tissues, capturing not only neural but also metabolic, apoptotic, and immune-related biological processes. By integrating two tissue types, the method naturally detects a broader spectrum of systemic pathways, offering mechanistic insight into processes underlying shared variance between brain regions and peripheral blood. The gene sets and pathways identified from the brain–blood PC intersections were then used as input for subsequent region-to-region analyses within the brain. We further prioritized pathways relevant to the olfactory region, extracting and interpreting those biological processes that showed shared variation detectable in both brain and blood.

## Results

To evaluate the influence of Alzheimer’s disease-related risk factors on olfactory-guided behavior, we assessed odor preference, anhedonia-like responses, and exploratory dynamics in mice stratified by APOE genotype, age, sex, diet, and immune background (hNOS2 vs. mNos2). Across behavioral assays, we observed consistent and converging evidence that both intrinsic (e.g., genotype, sex) and extrinsic (e.g., diet, age, immune status) factors interactively modulate olfactory behavior.

### Odor Salience

Odor salience (**Figure 2**), measured via normalized odor exploration across escalating concentrations of lemon odorant, showed a robust main effect of concentration (F(3, 1669) = 7.46, p < 0.001, η² = 0.0116, f = 0.108), indicating stimulus-dependent modulation of exploratory behavior. A significant concentration × age interaction (F(3, 1669) = 7.77, p < 0.001, η² = 0.0121) suggested age-related changes in odor tuning. Additional interactions with biological traits were significant, including concentration × diet (F(3, 1669) = 6.67, p < 0.001, η² = 0.0104), concentration × APOE × diet (F(6, 1669) = 3.26, p = 0.003, η² = 0.0102), and concentration × APOE × HN (F(6, 1669) = 2.22, p = 0.039, η² = 0.0069). A five-way interaction (concentration × APOE × age × HN × sex) also reached significance (F(6, 1669) = 3.86, p < 0.001, η² = 0.0120), highlighting a highly multifactorial regulation of odor-guided responses.

**Figure 2.**
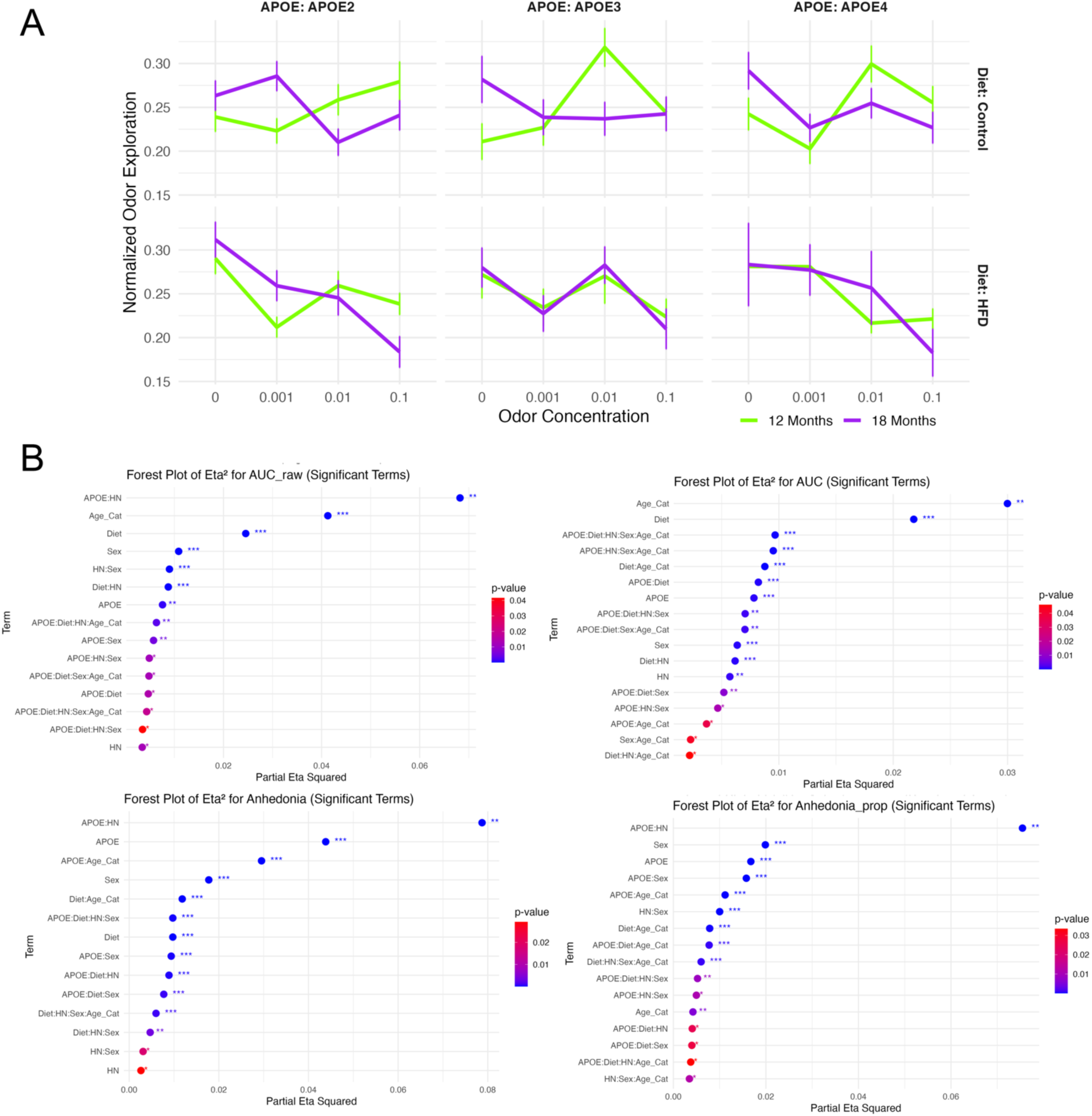
Odor salience is modulated by genotype, age, and diet. (A) Normalized odor exploration across four odor concentrations stratified by APOE genotype, age (12 vs. 18 months), and diet (control vs. high-fat). Each subplot represents group means ± SE. (B) Forest plots of partial eta squared (η²) effect sizes for significant terms (p < 0.05) in models assessing raw odor salience (AUC), normalized Anhedonia, and proportional Anhedonia. Terms are sorted by η², with significance stars indicating thresholds: \**p* < 0.05*, **p<0.01**, \*\*\***p<0.001*.

### Anhedonia

Anhedonia-like behavior, estimated from both total and proportional time exploring odorized vs. neutral stimuli, was significantly modulated by APOE × HN interactions. For absolute time, the interaction was highly significant (F(2, 1669) = 77.25, p < 0.001, η²_partial = 0.079, f = 0.292), with consistent effects in the proportional analysis (F(2, 1669) = 73.77, p < 0.001, η²_partial = 0.075). Main effects of APOE (F(2, 1669) = 41.46, p < 0.001, η²_partial = 0.044), sex (F(1, 1669) = 32.69, p < 0.001, η²_partial = 0.018), and diet × age (F(1, 1669) = 21.66, p < 0.001, η²_partial = 0.012) were also observed. Several higher-order interactions contributed to the behavioral variance, including APOE × sex, APOE × diet × HN, and diet × HN × sex × age (all p < 0.01). Together, these findings support the hypothesis that diminished reward-seeking under olfactory stimulation is shaped by genotype-immune interactions and modulated by age, sex, and diet.

### Exploratory Dynamics

Cumulative exploratory behavior quantified via area under the curve (AUC) metrics demonstrated robust effects of age and environment. For normalized AUC, age had the strongest main effect (F(1, 1669) = 56.02, p < 0.001, η²_partial = 0.030), followed by diet (F(1, 1669) = 40.38, p < 0.001, η²_partial = 0.022) and sex (F(1, 1669) = 11.60, p < 0.001, η²_partial = 0.0064). Diet × age interaction (F(1, 1669) = 16.04, p < 0.001) and higher-order interactions (e.g., APOE × diet × HN × sex × age, η²_partial = 0.0097; APOE × HN × sex × age, η²_partial = 0.0095) reflected the complexity of modulatory influences. These findings suggest that signal integration across concentrations (as reflected in AUC) is highly sensitive to the intersection of genotype, immune status, and developmental stage. Raw AUC analysis confirmed these patterns, with APOE × HN again emerging as the strongest interaction (F(2, 1669) = 66.32, p < 0.001, η²_partial = 0.068). Main effects of age (F(1, 1669) = 77.99, p < 0.001), diet (F(1, 1669) = 45.58, p < 0.001), and sex (F(1, 1669) = 19.90, p < 0.001) were robust, alongside immune-related interactions (HN × sex and diet × HN, both p < 0.001). Raw AUC thus provides a complementary metric of behavioral sensitivity to aging and immune factors, highlighting how environmental and genetic variables shape odor-guided behavior. These results suggest that the influence of APOE genotype on odor-preference and odor - guided behavior is significantly modulated by both diet and host background (HN), particularly in the context of aging. The high effect sizes for APOE × HN and Diet × Age_Cat interactions suggest compounded vulnerability in APOE4 mice on a high-fat diet.

### Habituation to Repeated Odor Exposure

We first analyzed total exploration time across all four objects during repeated odor presentations (Trials 1–4) and upon odor substitution (Trial 5) to assess general habituation (**Figure 3**).

**Figure 3.**
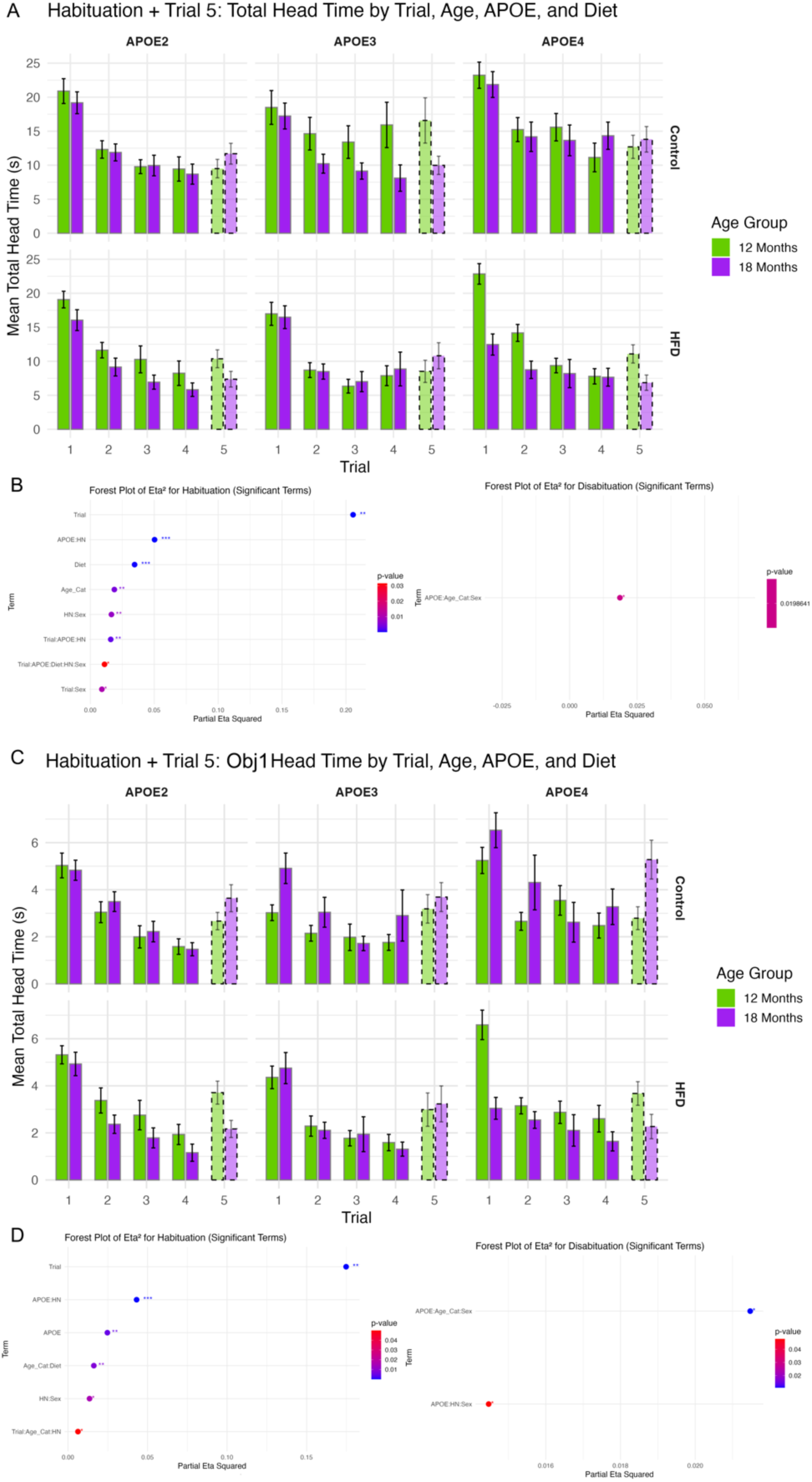
Effects of age, APOE genotype, and diet on olfactory habituation and dishabituation. **(A)** Total head time across all four objects declined over trials in all groups, consistent with habituation. APOE genotype, age (12 vs. 18 months), and high-fat diet modulated overall exploratory behavior, with older and HFD-fed mice showing reduced engagement. **(B)** Forest plot of significant terms in the mixed-effects model for total head time (habituation). Trial and multiple interactions involving APOE, HN, and diet contributed significantly. **(C)** Head time directed at the odor-switched object (Obj1) also showed trial-dependent habituation, but with stronger genotype-specific effects, particularly in APOE4 mice. **(D)** Forest plot of significant terms in the model for Obj1-directed head time (dishabituation). Significant three-way interactions involving APOE, age, sex, and housing suggest specific disruptions in novelty detection.

Across genotypes and diets, animals showed significant reduction in head time across trials (F(3,1240) = 106.94, p < 1.4e−61, η² = 0.21), consistent with robust habituation (**Figure 3A**). Diet (F(1,425) = 15.26, p < 0.0001, η² = 0.035) and age (F(1,425) = 8.14, p < 0.005, η² = 0.019) also contributed independently to exploratory behavior, with older and high-fat diet–fed mice generally showing reduced overall engagement.

To isolate responses to the odor-switched object (Obj1), we examined head time directed specifically at Obj1 across trials (**Figure 3B**). A similar pattern of habituation was observed (F(3,1241) = 87.70, p < 1.8e−51, η² = 0.175), though with additional significant effects of APOE genotype (F(2,425) = 5.39, p < 0.005, η² = 0.025), revealing genotype-specific modulation of odor investigation.

### Dishabituation to Novel Odor

Dishabituation was evaluated by comparing Trial 5 head time directed at Obj1 (odor-switched) versus prior trials. While no main effects of genotype, age, or diet reached significance when analyzing dishabituation responses across all four objects or for Obj1 alone, significant three-way interactions emerged. Specifically, APOE × Age × Sex and APOE × HN × Sex interactions reached significance (p = 0.02 and p = 0.04, respectively; **Figure 3C**), with modest effect sizes (η² ≈ 0.02). These findings suggest sex-specific modulation of dishabituation in the context of APOE genotype and housing conditions, potentially reflecting altered odor novelty detection in select subgroups.

### Odor Recognition Memory Across Delay Intervals

Recognition index (RI) scores were used to evaluate short-term (1h), long-term (24h), and remote (48h) odor memory across APOE genotypes, diets, and age groups (Figure 4A). While all groups displayed intact short-term memory (RI > 0.5), genotype and age-related differences emerged with longer delays. At 24 hours, APOE2 control-fed mice at 12 months showed the highest RI, with age-associated decline at 18 months. APOE3 and APOE4 groups exhibited flatter or diminished performance across age and diet conditions. Mixed effects modeling revealed significant interactions for APOE × HN, APOE × Diet × Age × Category, and Sex × HN (Figure 4B; partial η² = 0.028, p < .01), indicating sensitivity of long-term memory to both genetic and environmental modifiers. At 48 hours, all groups showed reduced recognition, with APOE × HN and APOE × Age interactions reaching statistical significance (partial η² = 0.032, p < .01), highlighting progressive deficits in odor memory consolidation and retrieval under risk-related conditions.

**Figure 4.**
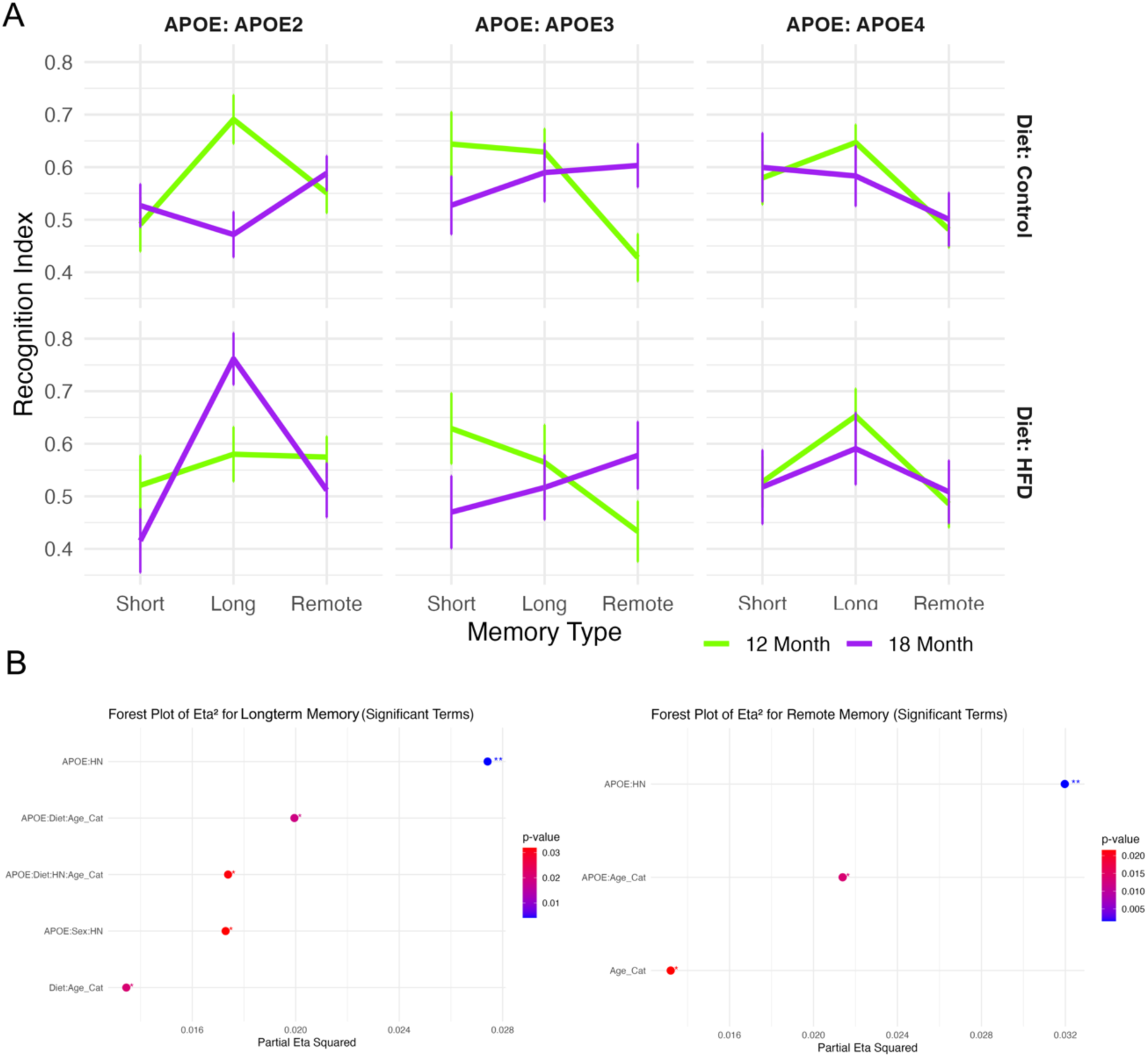
Odor recognition memory performance varies by APOE genotype, age, and diet across delay intervals. (A) Recognition Index (RI) was assessed after short-term (1 hour), long-term (24 hours), and remote (48 hours) delays. Across genotypes, animals showed intact short-term memory (RI > 0.5). Long-term memory showed age- and genotype-specific effects, with APOE2 mice on a control diet displaying elevated RI at 12 months and reduced RI at 18 months. APOE3 and APOE4 groups showed flatter trajectories with diet- and age-associated decline. Remote memory was impaired in all groups, especially with advancing age. (B) Forest plots show significant interaction effects on long-term and remote memory. Long-term recognition was significantly modulated by APOE × HN, APOE × Diet × Age, and Sex × HN interactions (partial η² up to 0.028). Remote memory showed significant contributions from APOE × Age and APOE × HN (partial η² up to 0.032).

For short-term memory, the linear model revealed no significant trait-specific differences in recognition index. In the long-term memory test, several significant interaction effects were observed, including Age × Diet [F(1, 395) = 5.4, p = 0.02], APOE × HN genotype [F(2, 395) = 5.6, p = 0.004], Age × APOE × Diet [F(2, 395) = 4.0, p = 0.02], and APOE × Sex × HN [F(2, 395) = 3.5, p = 0.03]. For remote memory, Age was a significant factor [F(1, 398) = 5.3, p = 0.02], along with significant interactions between Age × APOE [F(2, 398) = 4.3, p = 0.01] and APOE × HN [F(2, 398) = 6.6, p = 0.002]. In the linear mixed-effects model that included memory type as a stage factor, both Age [F(2, 1179) = 4.9, p = 0.008] and APOE [F(4, 1179) = 2.8, p = 0.03] were significant main effects. Additionally, significant interactions were observed for Age × APOE [F(4, 1179) = 3.4, p = 0.009] and APOE × HN [F(4, 1179) = 5.7, p = 0.0001]. Overall, Age and APOE emerged as consistent and significant factors across memory stages, and these effects—along with the Age × APOE interaction—are visualized in **Figure 4**.

The APOE2 genotype showed a lower recognition index in short-term memory but a higher recognition index in remote memory compared to other genotypes, as shown in **Figure 4A**. Age-related differences were more pronounced in **Figure 4B**, where younger mice exhibited higher recognition indices for short-term and long-term memory but lower indices in remote memory. To better visualize the interaction between APOE genotype and age across memory stages, we plotted memory trajectories in **Figure 4C**. All APOE groups followed a similar trend—recognition index increased from short-term to long-term memory, then declined in remote memory, though the extent of change varied. In the 12-month cohort, APOE2 mice showed the smallest drop in recognition index from long-term to remote memory, indicating better retention compared to APOE3 and APOE4. APOE3 mice had the steepest decline, followed by APOE4. In the 18-month cohort, APOE4 mice showed the smallest improvement from short-term to long-term memory and the largest decline from long-term to remote memory, suggesting greater memory impairment with age compared to APOE2 and APOE3.

### ROI Clusters Highlighting Olfactory Regions

An ROI analysis centered on 18 olfactory memory preselected regions revealed how these map in relation to each other, and the impact of averaged risk factors on image metrics including volume, diffusion and mean apparent propagator (**Figure 5**). The regions encompassed the olfactory system and associated limbic/memory circuits, including: Piriform Cortex, Amygdalopiriform Transition Area, Lateral Olfactory Tract, Ectorhinal Cortex, Caudomedial Entorhinal Cortex, Dorsal Intermediate Entorhinal Cortex, Dorsolateral Entorhinal Cortex, Medial Entorhinal Cortex, Amygdala (including Basolateral Amygdala), Accumbens, Claustrum, Cingulum, Ventral Orbital Cortex, Anterior Commissure, Bed Nucleus of the Stria Terminalis, and Ventral Tegmental Area. A hierarchical clustering based heatmap revealed that several clusters are enriched in olfactory regions, indicating shared effect size patterns across metrics in these structures. Clustering was based on Euclidean distance and complete linkage. Only ROIs with statistically significant effects (*FDR* < 0.05) were included. Together, these analyses show that (1) primary olfactory regions cluster together based on shared patterns of imaging changes under AD risk, and (2) the strongest risk-associated effects occur not only in traditional olfactory cortices but also in limbic and reward-related structures that integrate odor information with memory and motivational valence.

**Figure 5.**
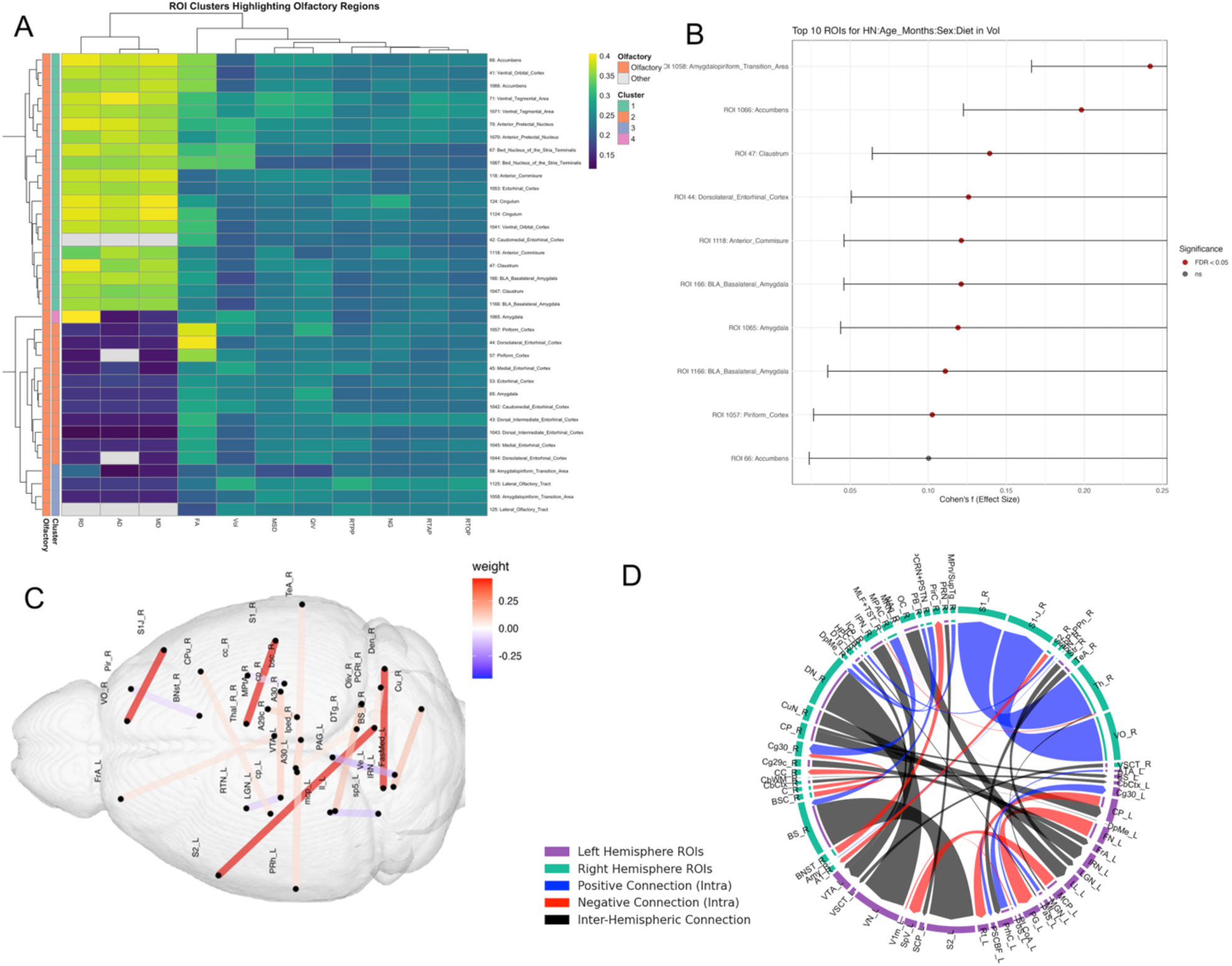
Clustering of olfactory-memory ROIs and top region effect sizes. (A) Hierarchical clustering of 18 a priori–selected olfactory–memory regions based on their Cohen’s f effect sizes across 11 imaging metrics (volume %, FA, MD, AD, RD, MSD, QIV, RTOP, RTAP, RTPP, NG). Branch lengths reflect Euclidean distance; complete linkage was used. The left color bar shows cluster membership (1–4), and the second bar flags true olfactory regions (orange) versus all others (gray). Note that cluster 2 (teal) is highly enriched for core olfactory areas (piriform cortex, amygdalopiriform transition area, anterior commissure, etc.), indicating these regions share a distinct signature of AD-risk effects on microstructure and volume. Cluster 4 (pink) captures more associative/hub regions (ventral orbital cortex, nucleus accumbens). Hierarchical clustering heatmap of brain regions (ROIs) based on average Cohen’s *f* effect sizes across multiple imaging-derived metrics. Each row represents a brain ROI, and each column corresponds to a metric, including fractional anisotropy (FA), radial diffusivity (RD), axial diffusivity (AD), mean diffusivity (MD), volume percentage (Vol), and advanced diffusion and microstructure metrics (MSD, QIV, RTPP, RTAP, RTOP, NG). ROIs are annotated by cluster membership (left bar) and olfactory categorization (second left bar), with olfactory ROIs marked in orange and others in light gray. (B) Forest-style dot plot of the top ten regions ranked by absolute Cohen’s f (random-effects meta-analysis across risk factors), showing point estimates for regional relative volumes (colored by olfactory vs. other) and 95% confidence intervals. The amygdalopiriform transition area and nucleus accumbens exhibit the largest effect sizes (f≈0.24 and 0.20, respectively), underscoring their vulnerability to combined APOE, diet, age, sex, and immune-background influences. Several canonical olfactory structures (piriform cortex, basolateral amygdala, anterior commissure) also rank among the top ten, highlighting that alterations in primary olfactory nodes parallel those in downstream reward and associative circuits. (C) Top 20 Elastic MCCA connectivity weights, shown in the bacground of the brain anatomy (D) Chord diagram showing the top connections and their weights follwing elastic net MCCA.

Elastic MCCA identified brain networks associated with risk factor traits including age, APOE genotype, diet, sex, and HN genotype as well as odor-based behavioral metrics. The sum of correlations was 1.267, with 1,000 bootstrap iterations yielding a 95% confidence interval of [1.183, 1.846] and a p-value < 0.0001. Following Elastic Net regularization, 242 connections were retained, which were further reduced to 39 through MCCA. Applying a threshold to exclude connections with weights below 0.05 resulted in a final set of 20 connections, as shown in **Figure 5**. Following Elastic Net filtering, the connectivity matrix retained non-zero weights for 49 connections. Of these, connections with absolute weights less than 0.05 were excluded, resulting in 20 significant connections reported in **Table 2**.

**Table 2.** Elastic MCCA selected connectivity list and associated weights for the top graph edges. The connections are ranked from top to bottom based on absolute weight values.

| Region 1 | Region 2 | Weight |
| --- | --- | --- |
| Right Ventral Orbital Cortex | Right Primary Somatosensory Cortex Jaw Region | 0.452 |
| Right Brain Stem | Left Secondary Somatosensory Cortex | 0.452 |
| Right Dentate Nucleus of Cerebellum | Left Vestibular Nuclei | 0.452 |
| Right Thalamus | Right Primary Somatosensory Cortex | 0.452 |
| Left Fastigial Medial Nucleus of Cerebellum | Left Periaqueductal Grey | -0.158 |
| Right Olivary Complex | Left Lateral Lemniscus | 0.144 |
| Right Cuneate Nucleus | Left Intermediate Reticular Nucleus | 0.143 |
| Right Interpeduncular Nucleus | Left Ventral Tegmental Area | 0.137 |
| Left Middle Cerebellar Peduncle | Left Spinal Trigeminal Nerve | -0.117 |
| Right Cerebral Peduncle | Left Cerebral Peduncle | 0.115 |
| Left Cerebral Peduncle | Left Reticular Nucleus of Thalamus | -0.103 |
| Right Medial Parietal Association Cortex | Right Cingulate Cortex | 0.101 |
| Left Perirhinal Cortex | Left Cingulate Cortex | 0.0837 |
| Right Bed Nucleus of the Stria Terminalis | Right Piriform Cortex | -0.0717 |
| Right Cingulate Cortex | Left Frontal Association Cortex | 0.0684 |
| Right Striatum | Left Lateral Geniculate Nucleus | 0.0641 |
| Right Temporal Association Cortex | Left Perirhinal Cortex | 0.0630 |
| Right Parvocellular Reticular Nucleus and Principal Sensory Trigeminal Nucleus | Right Brain Stem | 0.0576 |
| Right Brachium of Superior Colliculus | Right Corpus Callosum | -0.0528 |
| Right Dorsal Tegmentum | Left Ventral Tegmental Area | 0.0519 |

The strongest weighted connections (weight = 0.452) included: right brainstem to left secondary somatosensory cortex, right dentate nucleus of the cerebellum to left vestibular nuclei, right thalamus to right primary somatosensory cortex, and right ventral orbital cortex to right primary somatosensory cortex. The next highest absolute weight was observed between the left fastigial medial nucleus of the cerebellum and the left periaqueductal gray (weight = -0.158). As shown in **Figure 5**, the presence of both positive and negative weights suggests a combination of strengthened and compensatory connections associated with increased scent preference and novel odor recognition index.

### Gene Module and Microstructure Coupling

To investigate whether peripheral markers of gene expression reflect changes in in olfactory-memory circuits, we first performed transcriptomic analyses from whole blood RNA using principal component analysis (PCA), which identified 10 modules which we tested for co-expressed gene clusters. Functional enrichment analysis revealed several modules that were biologically meaningful, particularly in the context of olfaction and memory. As shown in **Table 3**, Eigengenes 1 to 3 showed the most prominent enrichment for biologically relevant pathways. Eigengene 1 was significantly associated with pattern recognition receptor signaling (p = 0.003), suggesting involvement in early sensory detection and immune-like receptor activity. Eigengene 2 was enriched for phagocytosis and recognition-related pathways (p = 0.007), implicating roles in sensory clearance and responsiveness to external stimuli. Notably, Eigengene 3 showed strong enrichment for learning and memory functions (p = 0.002), including processes involved in hippocampal development—highlighting a direct link between gene regulation and cognitive mechanisms underlying olfactory memory formation. Eigengenes 4 through 8 demonstrated moderate enrichment for pathways such as neuron recognition, cell-cell signaling, and receptor activity. For example, Eigengene 4 was associated with neuron recognition (p = 0.010), while Eigengene 8 showed enrichment for cell adhesion involved in neuronal migration (p = 0.029), suggesting supportive roles in neuronal connectivity and communication. In contrast, Eigengenes 9 and 10 did not show statistically significant enrichment for olfactory or memory-related terms after correction, indicating that they may reflect unrelated or background biological variation within the transcriptomic dataset.

**Table 3.** Top enriched biological pathways associated with eigengene modules for blood. Modules 1 to 3 show strong relevance to sensory recognition, 4 to 8 show neuronal signal and development and modules 9 and 10 did not yield significant enrichment related processes.

| Eigengene Module | Top Enriched Pathway | P-value | Notes |
| --- | --- | --- | --- |
| Eigengene 1 | Pattern recognition receptor signaling | 0.003 | Strong immune/sensory relevance |
| Eigengene 2 | Phagocytosis, recognition | 0.007 | Sensory clearance processes |
| Eigengene 3 | Learning or memory | 0.002 | Direct cognitive link (memory formation) |
| Eigengene 4 | Neuron recognition | 0.010 | Supports neuronal differentiation |
| Eigengene 5 | Pattern recognition receptor activity | 0.015 | Supports sensory input mechanisms |
| Eigengene 6 | Cell surface receptor signaling | 0.018 | Moderate sensory/cognitive relevance |
| Eigengene 7 | Cell-cell recognition | 0.022 | Intercellular signaling |
| Eigengene 8 | Cell adhesion involved in neuron migration | 0.029 | Neuronal development process |
| Eigengene 9 | — | — | No significant olfactory/memory pathways |
| Eigengene 10 | — | — | Minor enrichment, not reaching significance |

To assess whether specific gene expression modules covary with local brain structure, we examined correlations between blood-derived eigengenes and regional microstructural MRI metrics across eight predefined olfactory-memory regions. After applying FDR correction, we identified 431 significant eigengene– region correlations (FDR < 0.05), highlighting widespread and spatially specific molecular–structural coupling within these circuits. The strongest correlations were observed in the Parasubiculum, where diffusion-based metrics—including axial diffusivity (AD), radial diffusivity (RD), and mean diffusivity (MD)—were tightly linked to Eigengene 2 (**Table 4**). The top five absolute correlations (|r| ≈ 0.48–0.49) all involved Eigengene 2 and were highly significant (p_FDR < 1 × 10^-30^), with R² values indicating that up to 24% of the variance in regional diffusivity could be explained by this eigengene. Notably, similar associations were found in the Hippocampus and Postsubiculum and Piriform Cortex, underscoring the relevance of Eigengene 2 to microstructural variation across the subicular–hippocampal axis.

**Table 4.** Top region-eigengene correlation between diffusion MRI metrics and transcriptomic modules in olfactory-memory related structures demonstrate a role for memory and olfaction related regions.

| Structure | Metric | Eigengene | r | p_FDR | R<sup>2</sup> |
| --- | --- | --- | --- | --- | --- |
| Parasubiculum | AD | Eigengene_2 | −0.4919 | $1.6 \times 10^{-32}$ | 0.24 |
| Parasubiculum | MD | Eigengene_2 | −0.4870 | $8.6 \times 10^{-32}$ | 0.24 |
| Parasubiculum | RD | Eigengene_2 | −0.4836 | $3.2 \times 10^{-31}$ | 0.23 |
| Hippocampus | AD | Eigengene_2 | −0.4780 | $7.7 \times 10^{-31}$ | 0.23 |
| Postsubiculum | AD | Eigengene_2 | −0.4777 | $8.1 \times 10^{-31}$ | 0.23 |
| Piriform Cortex | MSD | Eigengene_8 | 0.3636 | $2.1 \times 10^{-12}$ | 0.14 |
| Piriform Cortex | FA | Eigengene_2 | 0.3176 | $6.8 \times 10^{-13}$ | 0.10 |
| Piriform Cortex | NG | Eigengene_5 | −0.3237 | $2.8 \times 10^{-11}$ | 0.13 |
| Piriform Cortex | RTOP | Eigengene_8 | −0.3102 | $2.4 \times 10^{-9}$ | 0.10 |
| Piriform Cortex | RTOP | Eigengene_4 | −0.3089 | $5.9 \times 10^{-11}$ | 0.12 |
| Hippocampus | MD | Eigengene_2 | −0.4758 | $1.5 \times 10^{-30}$ | 0.23 |
| Hippocampus | RD | Eigengene_2 | −0.4738 | $2.8 \times 10^{-30}$ | 0.22 |
| Hippocampus | FA | Eigengene_2 | 0.3689 | $2.7 \times 10^{-17}$ | 0.14 |
| Hippocampus | QIV | Eigengene_8 | 0.3643 | $2.1 \times 10^{-12}$ | 0.14 |

Eigengene 2 emerged as a dominant driver of structural variability within deep olfactory-memory regions, particularly those implicated in memory encoding and contextual sensory integration. In contrast, other eigengene modules demonstrated more regionally restricted associations. For instance, Eigengene 3 showed moderate correlations (r ≈ 0.3–0.4) with diffusion metrics in the Piriform Cortex, while Eigengene 4 was preferentially associated with the Ventral Orbital Cortex. These results suggest modular specificity in how transcriptomic variation relates to local brain architecture across the olfactory-memory system. Interestingly, volume-based metrics (e.g., regional volume percentage) yielded much weaker associations overall (|r| < 0.3), suggesting that diffusion MRI is more sensitive than regional volumetry in capturing molecular influences on tissue structure. This observation supports the utility of advanced diffusion metrics in detecting subtle gene-related variability in brain architecture.

Thus, these findings suggest that Eigengene 2 captures molecular processes, potentially related to lipid metabolism, myelination, or neuroinflammatory signaling, tightly coupled to microstructural integrity in key olfactory-memory hubs such as the subiculum and hippocampus. Modules such as Eigengenes 3, 4, 5, and 8 appear to mediate structural variation in more anterior olfactory regions, including the Piriform and Ventral Orbital Cortices. The breadth of significant eigengene–MRI associations reflects both the biological richness of the blood transcriptome and the sensitivity of diffusion imaging to detect microstructural changes relevant to olfactory behavior and memory.

Across modules, the top loading genes highlight a rich interplay between synaptic plasticity (Grin2a, Camk2a, Snap25), metabolic support (Cs, Cox5a, Uqcrc2), glial modulation (Gfap, S100b), and myelination (Plp1, Mog). Multiple genes (e.g., Nptx2, Camk2a, Grin2a) have well-established links to both memory and olfactory processing, suggesting these eigengene modules capture biologically relevant axes of neural function underlying sensory memory and circuit plasticity.

### Mapping Peripheral to Central Transcriptomic Processes

To elucidate the molecular coupling between the brain and peripheral blood, and then more focused on the olfactory region and other brain regions, we performed principal component analysis (PCA) on transcriptomic data from each compartment (**Figure 6A–B**). For each region, the top four principal components (PCs) were computed, and all pairwise correlations between Olf PCs and those from other regions or blood samples were assessed, with statistical significance determined using the false discovery rate (FDR).

**Figure 6.**
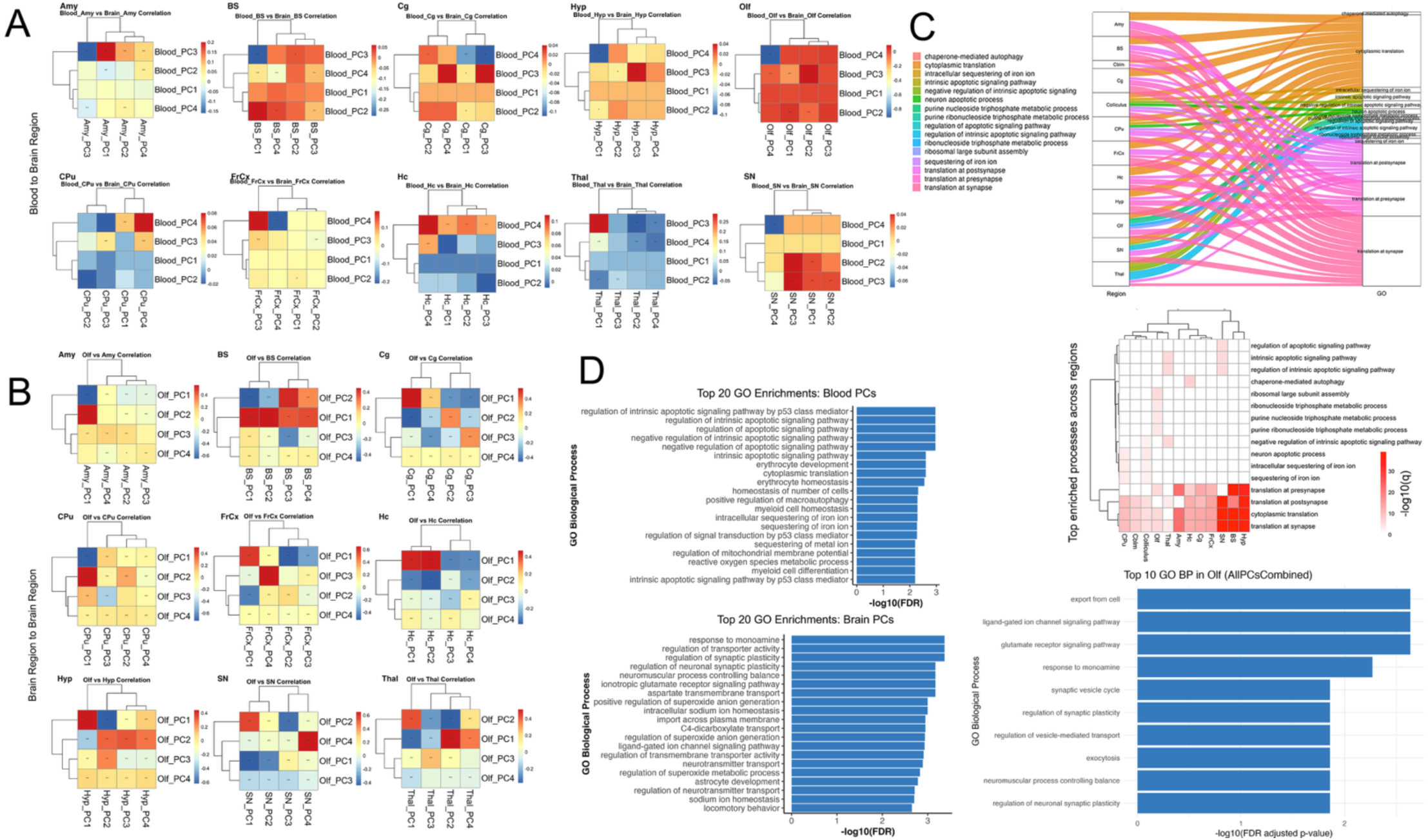
Mapping Peripheral to Central Transcriptomic Processes. (A) Heatmaps illustrating the correlation between blood-derived principal components (PCs) and brain PCs. Abbreviations: Hc, hippocampus; Olf, olfactory bulbs; FrCx, frontal cortex; CPu, caudate putamen; Hyp, hypothalamus; BS, brainstem; SN, substantia nigra; Cg, cingulate cortex; Amy, amygdala; Thal, thalamus; Colliculus, superior and inferior colliculi; Cbls, cerebellum. *p < 0.05. (B) Heatmaps illustrating the correlation between olfactory region–derived principal components (PCs) and PCs from other brain regions. *p < 0.05; **p<0.01; *** p<0.001. (C) Sankey diagram depicting pathways derived from genes driving enriched pathways for significantly correlated PCs between brain regions and blood. Right: clustered heatmap showing convergence of blood to region-specificity of enriched biological processes. (D) Barplots showing the top enriched biological processes for blood PCs associated with brain regions, brain region–to–brain region; and olfactory region analyses. Bars represent –log10(FDR) for each process.

### Principal Component Correlations Reveal Regional and Brain–Blood Transcriptomic Coupling

We identified numerous significant correlations (FDR < 0.05) between specific Olf PCs and PCs from other brain regions (**Figure 6B**), as well as between Olf PCs in blood and PCs from other regional blood samples (Fig. 6A). The strongest associations were found for olfactory–hippocampus and olfactory–thalamus PC pairs, with both positive and negative correlations observed across principal axes. These patterns indicate that only a subset of major transcriptomic axes underlie inter-regional and brain–blood coupling, reflecting targeted, rather than global, coordination of gene expression programs.

### Top Genes and Pathways in Blood–Brain and Brain–Brain Coupling

For each significant PC pair, we identified the top genes by absolute PC loading that contributed most to the observed correlations. Gene ontology enrichment analysis of these high-loading genes revealed strong convergence on pathways involved in apoptotic signaling, erythrocyte development, oxidative stress response, synaptic plasticity, and transporter activity.

In blood, several canonical regulators of oxidative defense and metabolism—including Gpx1, Sod2, Nqo1, and Slc2a1—were among the most highly loaded genes in significant PCs, underscoring the central role of redox and metabolic homeostasis in mediating peripheral transcriptomic signals associated with brain states. Notably, blood PCs were also enriched for pathways related to synaptic plasticity, neurotransmitter signaling, and immune function, suggesting that blood transcriptomic profiles can reflect key aspects of central nervous system activity and vulnerability.

Within the brain, significant PC pairs were predominantly driven by high-impact genes involved in apoptosis (Trp53, Casp3, Bax), synaptic regulation (Grin2b, Dlg4, Syn1), and metabolic stress (Sod1, Cat, Gpx1, Nrf2, Hmox1). The most significantly enriched biological processes included intrinsic apoptotic signaling (notably p53-mediated), cytoplasmic translation, synaptic transmission, and erythrocyte homeostasis. Several genes— including Gpx1, Nqo1, and Slc2a1—appeared in both blood and brain analyses, highlighting systemic coordination and the potential for shared molecular mechanisms.

### Pathway Coupling and Enrichment Analysis

Pathway coupling was illustrated by Sankey diagrams and a heatmap (**Figure 6C, top**), demonstrating that processes such as apoptosis, protein translation, synaptic function, and metabolic regulation were consistently represented among the top genes across multiple regions. The heatmap confirmed that these pathways were enriched in several, but not all, region pairs, reflecting both global and region-specific molecular drivers.

Across the most significant blood–brain principal component pairs for the olfactory region, we observed strong enrichment for pathways involved in protein synthesis, metabolic regulation, and cellular transport. Specifically, the leading processes included cytoplasmic translation and translation at the presynapse, ribosomal large subunit biogenesis, carbohydrate derivative catabolic process, purine-containing compound metabolic process, and protein localization to cell–cell junctions. These findings highlight the shared regulation of translation and synaptic protein synthesis, as well as core metabolic pathways, between blood and olfactory brain tissue. The presence of these intersecting pathways suggests a robust molecular link between peripheral and central compartments, emphasizing processes essential to olfactory neuronal function and supporting the idea that blood-based gene expression can reflect key brain-region vulnerabilities (Figure 6C bottom).

To provide a comprehensive perspective, we visualized the top 20 significantly enriched GO biological processes for both brain and blood (**Figure 6D**). In the brain, intrinsic apoptotic signaling, cytoplasmic translation, erythrocyte development and homeostasis, and synaptic plasticity were among the most enriched pathways. In blood, enrichment included apoptotic processes, canonical neural pathways (regulation of synaptic plasticity, ionotropic glutamate receptor signaling, monoamine response), transporter activity, immune processes, and oxidative stress responses. Several pathways—such as oxidative stress response, iron ion sequestration, and metabolic homeostasis—were common to both compartments, underscoring their centrality in coordinated molecular responses.

Collectively, these results demonstrate that transcriptomic coupling between olfactory, other brain regions, and blood is driven by a core set of high-impact genes acting within key biological pathways, including apoptosis, synaptic regulation, metabolic, and oxidative stress responses. The broader view provided by the top 20 enrichments further underscores the diversity and complexity of the processes mediating brain–blood communication and regional brain coordination. The convergence of these processes across compartments supports a model of integrated molecular signaling underpinning both regional brain interactions and brain– blood communication.

### Enrichment of Olfactory Synaptic Pathways Among Genes Shared Between Brain and Blood

To further dissect the molecular mechanisms linking brain and blood, we prioritized pathways and genes relevant to the olfactory region, focusing on biological processes that exhibited significant shared variation between olfactory brain tissue and blood gene expression. Gene Ontology enrichment analysis of the top intersecting genes revealed robust enrichment for pathways essential to olfactory function, most notably the glutamate receptor signaling pathway and ligand-gated ion channel signaling pathway (FDR-adjusted p = 0.0023). These pathways are fundamental for synaptic transmission within olfactory circuits, where glutamatergic signaling is central to conveying odor information from olfactory sensory neurons to the olfactory bulb and higher-order brain regions. In addition to glutamatergic signaling, we found significant enrichment for biological processes such as regulation of synaptic plasticity, intracellular sodium and potassium ion homeostasis, and regulation of neurotransmitter transport. These mechanisms are critical for olfactory signal transduction and support the adaptive capacity of olfactory networks during odor learning and memory. The enrichment of these terms suggests that the shared molecular signatures between brain and blood are rooted in processes fundamental to olfactory system function. Several of the top shared genes are directly implicated in olfactory neuronal function and synaptic physiology. For example, Atp1a3 maintains neuronal ion gradients necessary for action potential propagation within olfactory pathways [40], while App is involved in synaptic organization and function [41] and is expressed in olfactory regions. Plp1 supports myelination and rapid signal conduction in the olfactory system [42], and Cplx2 regulates neurotransmitter release at olfactory synapses [43].

These results highlight a robust molecular bridge between peripheral blood and olfactory brain regions, implicating synaptic signaling, ion homeostasis, and neurotransmitter regulation as shared processes detectable across tissues. Our findings reinforce the central role of glutamatergic and synaptic pathways in olfactory function and demonstrate that blood-based molecular signatures can reflect region-specific brain vulnerabilities relevant to olfaction.

## Discussion

We examined how multiple AD risk factors—including age, APOE genotype, diet, sex, and HN genotype— influence odor preference, habituation/dishabituation, and memory performance in mice, and how these behavioral outcomes relate to underlying brain connectivity. Our findings show that each risk factor exerts distinct, and often interactive, effects on odor-guided and memory-related behaviors, highlighting the complexity of sensory-cognitive processes in the aging brain. One of our most striking findings was that different odorant (lemon) concentrations elicited distinct patterns across risk factors. Older mice generally showed higher preference for lower odor concentrations, while younger mice were more responsive to higher concentrations. These patterns may reflect age-related shifts in olfactory sensitivity—possibly due to reduced peripheral or central olfactory function in older animals, making higher concentrations aversive or overwhelming. Alternatively, motivational or exploratory differences could contribute, with older mice engaging more with weaker stimuli and younger mice exhibiting greater novelty-seeking behavior at stronger concentrations [44]. Additionally, prior studies have shown that mice prefer lemon over certain other scents, such as rose, and tend to spend more time digging in lemon-scented bedding— further supporting its salience as a behaviorally relevant odor [45].

APOE genotype effects were also evident in the habituation/dishabituation test, offering additional insight into how different genotypes adapt to repeated stimuli and respond to novelty. During the habituation phase, particularly in the final trial, APOE2 mice exhibited the lowest investigation time, suggesting more rapid adaptation. In the dishabituation trial, APOE2 mice showed higher dishabituation indices compared to APOE3 and APOE4 mice, indicating stronger re-engagement with the novel odor. APOE2 mice also displayed higher dishabituation indices compared to other genotypes for both control and high-fat diet (HFD) cohorts. Notably, the difference between APOE2 and other genotypes, especially APOE4, became more pronounced under HFD conditions. This suggests that while APOE4 mice may be more sensitive to the detrimental effects of HFD, APOE2 mice could exhibit compensatory or protective responses [46, 47]. These findings further support the concept of genotype-specific dietary vulnerabilities or resilience, reinforcing the potential for targeted dietary interventions to mitigate or delay sensory-cognitive decline in genetically at-risk populations [48]. Also, our results suggest that APOE2 mice not only habituate more efficiently but also sustain a strong novelty response—traits that may reflect underlying neuroprotective mechanisms, which aligns with previous research indicating that APOE2 confers resilience against neurodegenerative processes [47, 49]. In contrast, APOE4 mice, who carry the highest genetic risk for AD and age-related neurodegeneration, exhibit blunted novelty responses or delayed habituation, potentially reflecting reduced synaptic plasticity and impaired sensory processing associated with their genetic vulnerability [50, 51].

Memory performance measured across short-term, long-term, and remote time points provided further evidence of how genotype and age differentially influence memory consolidation and retrieval. Younger mice generally outperformed older mice in short-term and long-term memory tasks but showed reduced performance in remote memory tests. While it is typical for younger animals to excel during initial encoding and consolidation phases, the divergence observed at remote memory may reflect trade-offs or compensatory mechanisms in older cohorts [52–54]. Interestingly, APOE2 mice exhibited lower short-term memory performance but higher remote memory retention with aging compared to other genotypes. This pattern suggests that memory formation, consolidation, and retrieval may rely on partially distinct neural circuits whose efficiency varies by genotype and with age [47]. These findings highlight the need for further investigation such as using electrophysiological or imaging methods to better understand how hippocampal-to-cortical transitions during remote memory processing are shaped by APOE status and aging.

Alongside the behavioral findings, Elastic MCCA analyses identified key brain connections with both positive and negative weights. Positive connectivity weights may reflect stronger coupling that enhances odor recognition or memory, while negative weights could represent inhibitory or compensatory pathways that reorganize in response to risk factors such as age, diet, or genotype [55]. One notable connection is between the right ventral orbital cortex and the right primary somatosensory cortex, which exhibited one of the highest weights (0.452). The ventral orbital cortex is known for its role in assigning reward value to odors, integrating olfactory information with other sensory cues to guide behavior [56]. The primary somatosensory cortex, while traditionally associated with processing touch, temperature, and proprioception, has also been implicated in aspects of odor perception and cross-modal integration [57, 58]. Together, these regions may support the formation and retrieval of associations between olfactory stimuli, spatial context, and the motor components of exploratory behavior, such as approach, avoidance, and investigation [59].

In addition to the ventral orbital cortex–primary somatosensory cortex pathway, several other connections emerged that further highlight the interplay among olfactory, emotional, and reward circuits. First, the bed nucleus of the stria terminalis is an extended amygdala structure crucial for regulating stress and anxiety [60], while the piriform cortex is a primary olfactory region involved in odor discrimination and memory formation [61]. The negative correlation (–0.072) between these areas suggests that as piriform cortex function or connectivity declines, stress pathways regulated by the bed nucleus of the stria terminalis may become disinhibited, linking diminished olfactory processing to heightened emotional reactivity [62]. Another notable connection is between the fastigial medial nucleus of the cerebellum and the periaqueductal grey (–0.158). The fastigial medial nucleus, one of the deep cerebellar nuclei, helps coordinate motor and affective processes [63], whereas the periaqueductal grey is central to defensive and autonomic responses [64]. This connection has previously been studied *in vivo*, and the negative correlation could indicate that as the cerebellar nucleus modulates sensorimotor and emotional integration, periaqueductal grey–mediated defensive reactivity is reduced, or vice versa, reflecting changes in autonomic and visceral responses to odors [65]. Lastly, the interpeduncular nucleus and the ventral tegmental area demonstrated a positive correlation (0.137). The interpeduncular nucleus, located in the midbrain, has been implicated in reward processing, mood regulation, and certain olfactory-driven behaviors [66, 67], while the ventral tegmental area is a dopaminergic hub integral to motivation and reward [68]. This connection has been confirmed by viral tracing and these regions undergo synchronized activation for encoding motivated exploration which may explain that it potentially increases motivational salience or attentional focus on novel or preferred scents. [69].

Furthermore, these findings underscore how age, APOE genotype, and diet collectively shape olfactory preference, novelty detection, and memory processes in mice. The fact that APOE2 animals remain relatively resilient under a high-fat diet aligns with literature suggesting this genotype confers protection against metabolic and neurodegenerative stressors [47, 49]. By pinpointing distinct neural circuits linked to these behaviors—particularly connections involving the ventral orbital cortex and primary somatosensory cortex—our study highlights potential targets for future interventions, such as genotype-informed dietary strategies and therapies aimed at sustaining critical brain networks. Nevertheless, there are several limitations to pay attention to, including baseline exploratory drive, potential stress from handling, circadian factors, and variations in odor delivery especially in memory test, all of which could affect the quantitative metrics collected. Replication in independent cohorts would further establish the robustness of these findings, and more direct measures of neural activity (e.g., fMRI or in vivo electrophysiology) during odor tasks could validate the connectivity patterns reported here [70, 71]. From a translational perspective, extending these results to human populations, especially those carrying different APOE alleles or at risk from dietary factors, may help identify early biomarkers and interventions for age-related cognitive decline [72]. Even so, by integrating odor-based behavioral measures and network-level connectivity, our multifaceted approach offers a deeper understanding of how risk factors and neural connectivity markers converge to shape odor-based cognitive decline.

Our investigation of blood based eigengenes revealed 8 significant modules, enriched for genes central to synaptic transmission and plasticity, epigenetic and metabolic regulation, glial support, and myelination. Multiple modules highlight genes essential for learning and memory (*Camk2a*, *Grin2a*, *Snap25*, *Ppp1r1b*) and for olfactory circuit function (*Apoe*, *Gfap*, *Snap25*), supporting their roles in integrating sensory and cognitive processes. Principal components 1 through 8 (PC1–PC8) captured major axes of molecular variation associated with imaging properties of brain regions, each with distinct functional relevance to memory and olfactory circuits. PC1 was driven by genes related to chromatin structure and vesicle trafficking (e.g., *Nsd1*, *Wdr82*, *Vps35*), which are essential for neuronal gene regulation and synaptic remodeling underlying learning and memory [73]. PC2 featured core regulators of synaptic plasticity and signaling such as *Nptx2* and *Camk2a*, directly implicated in olfactory learning and spatial memory [74]. PC3 prioritized genes governing neurotransmission, especially synaptic vesicle machinery and NMDA receptor function (e.g., *Grin2a*, *Snap25*), fundamental for long-term potentiation and memory encoding [75] [76]. PC4 highlighted axon guidance and reward pathway genes (*Dpysl2*, *Fyn*, *Ppp1r1b/DARPP-32*), with key roles in dopaminergic signaling and memory circuits [77]. PC5 was enriched for ribosomal and elongation factor genes, supporting the protein synthesis required for memory consolidation and synaptic plasticity in both olfactory and hippocampal systems [78]. PC6 was dominated by mitochondrial electron transport genes (e.g., *Uqcrc2*, *Ndufa1*, *Cox5a*), reflecting neuronal metabolic needs during high-demand cognitive and sensory processing [79]. PC7 was defined by glial and lipid metabolism genes (*Gfap*, *Apoe*, *S100b*), highlighting the importance of astrocytic support and lipid regulation in both synaptic plasticity and olfactory function [80]. Finally, PC8 captured myelination and oligodendrocyte-associated transcripts (e.g., *Plp1*, *Mobp*), crucial for rapid axonal conduction and effective transmission of memory and sensory information [81]. Together, these modules illuminate the tightly coordinated molecular programs underlying memory and olfaction in the mammalian brain.

In this study, we identify significant associations between blood gene expression profiles related to actin cytoskeletal regulation and regional diffusion MRI metrics of brain microstructure. Pathways related to actin filament organization, actin polymerization, intracellular transport, and cytoskeletal remodeling were enriched across multiple eigengene modules, notably Eigengene 1, 2, 3, and to a lesser extent, 6 and 8. These findings suggest that peripheral regulation of cytoskeletal dynamics may reflect or influence microstructural brain properties detectable by diffusion imaging.

The actin cytoskeleton is essential for maintaining cellular structure and facilitating dynamic processes such as membrane extension and compaction during myelination. Prior studies have shown that actin polymerization promotes the initial expansion of oligodendrocyte membranes, while subsequent actin depolymerization is required for myelin sheath compaction[82, 83]. Our findings extend this knowledge by demonstrating that actin-related gene expression in the blood correlates with imaging-derived measures of brain tissue organization, implying a potential systemic signature of brain microstructural health.

One possible interpretation is that blood-based actin pathway expression mirrors ongoing cytoskeletal remodeling processes within the central nervous system, either through shared developmental programs, immune-mediated mechanisms, or vascular influences. Alternatively, actin dysregulation in the periphery may contribute to neuroinflammatory cascades that secondarily impact brain structure.

These results provide a foundation for exploring blood actin pathway markers as accessible indicators of brain health, particularly in aging and neurodegenerative contexts where cytoskeletal instability and myelin degradation are prominent features. Future studies integrating longitudinal blood sampling, multimodal imaging, and direct measures of myelin integrity will be crucial to determine causality and therapeutic potential.

A limitation of this study is that the associations are correlative and peripheral blood expression may not fully capture the complexity of brain-specific actin dynamics. Nevertheless, the convergence across multiple eigengene modules strengthens the biological plausibility of our findings.

We leveraged a principal component–based correlation framework to dissect the molecular relationships between the olfactory region, other brain regions, and peripheral blood. Our findings reveal that a small subset of principal components—those capturing the most salient gene expression variance—are strongly correlated across brain regions and between brain and blood, highlighting targeted, rather than global, coordination of transcriptomic programs. Notably, these correlations were both positive and negative, reflecting dynamic and sometimes antagonistic regulation of gene expression across neuroanatomical boundaries. We found that a core group of high-impact genes drives both region-to-region and brain–blood correlations, supporting the concept that key biological processes such as apoptosis, metabolic and oxidative stress response, and synaptic plasticity are central to neural system resilience and disease risk.

Gene ontology enrichment analysis demonstrated that the genes driving these correlated principal components converge onto core biological processes, with remarkable consistency between brain and blood. Our shared blood-brain prominent pathways included intrinsic apoptotic signaling (notably p53-mediated), cytoplasmic translation, ribosomal assembly, and erythrocyte homeostasis. The repeated enrichment of erythrocyte-related processes suggests an underappreciated neurovascular or systemic metabolic component to brain transcriptomic coordination, consistent with emerging evidence linking vascular and metabolic dysfunction to neurodegenerative disease. In the pool of shared brain region to brain regions pathways, we observed enrichment not only for apoptotic and metabolic pathways, but also for processes associated with synaptic plasticity, glutamate receptor signaling, and monoamine response, supporting the notion that peripheral blood transcriptomes can capture echoes of central nervous system state.

At the gene level, top-loading genes within these significant PCs include canonical mediators of apoptosis (Trp53, Bax, Casp3), synaptic function (Grin2b, Dlg4, Syn1), ribosomal and translational activity, and metabolic/oxidative stress responses (Sod1, Cat, Gpx1, Nrf2, Hmox1). The high-loading genes associated with metabolic and oxidative stress—such as Gpx1, Sod2, Nqo1, and Slc2a1—underscore the contribution of redox and energy homeostasis to brain–blood transcriptomic coupling and may represent critical nodes for cross-tissue signaling and biomarker development. The presence of neuron-centric pathways and genes in blood further supports the potential of blood-based omics as accessible biomarkers for brain health and disease.

The use of PCA enabled us to disentangle both dominant and subtle axes of transcriptomic variation, capturing synergistic as well as antagonistic relationships between regions and compartments. Our results support an integrated model in which coordinated regulation of apoptosis, metabolism, and synaptic plasticity underpins neural system resilience and vulnerability to disease.

Limitations of this study include the potential for residual confounding due to technical variation or differences in cellular composition. Moreover the PCA approach may under-represent rare but functionally important cell-type–specific signals. Nevertheless, the consistency of key pathways and genes across compartments, and across analytic approaches, strengthens the robustness of our findings.

In summary, our integrative transcriptomic analysis identifies conserved and coordinated molecular programs underpinning brain–blood communication and regional brain interactions. The central roles of apoptotic, metabolic, and synaptic pathways provide new insight into the mechanisms of neural resilience and vulnerability, and highlight promising targets for biomarker discovery and intervention in neurodegenerative disease.

Our integrated analysis of principal components (PCs) of gene expression across olfactory and other brain regions, as well as peripheral blood, reveals several key themes regarding the molecular pathways underpinning regional and systemic transcriptomic coupling. By focusing on the top-loading genes from significantly correlated PC pairs, we have delineated biological processes that may drive coordinated changes between the olfactory system and broader neural circuits, as well as between central and peripheral compartments.

In brain, enrichment analysis of these key gene sets consistently highlighted pathways involved in intrinsic apoptotic signaling, particularly those mediated by p53, as well as erythrocyte development, cytoplasmic translation, and homeostatic processes. The strong prominence of apoptotic pathways is consistent with extensive literature implicating programmed cell death and p53 signaling in the pathogenesis of Alzheimer’s disease and related neurodegenerative conditions. The observed enrichment for erythrocyte and homeostasis-related processes may reflect the growing appreciation of neurovascular and metabolic contributions to brain aging and cognitive decline. Our findings thus reinforce the concept that cell survival, maintenance of cellular homeostasis, and the integrity of neurovascular units are tightly interwoven in shaping transcriptomic states associated with disease vulnerability.

Perhaps more unexpectedly, analysis of blood-based PC correlations revealed enrichment for synaptic and neurotransmission-related pathways, including regulation of synaptic plasticity, ionotropic glutamate receptor signaling, monoamine response, and transporter activity. The detection of these canonical neuronal processes in peripheral blood may be attributable to several mechanisms, such as cross-talk between the immune and nervous systems, the presence of circulating neuronal vesicles, or systemic responses that parallel central changes. Moreover, oxidative stress and ion homeostasis pathways emerged in the blood signature, further supporting a model wherein peripheral molecular profiles can serve as accessible surrogates of central nervous system activity and health. These results align with and extend previous studies suggesting that blood transcriptomics can reflect aspects of brain pathology, particularly when leveraging dimension reduction approaches that capture coordinated variance [84]. The convergence of apoptotic, synaptic, and metabolic stress pathways across central and peripheral compartments underscores the complexity of neurodegenerative disease biology and highlights the utility of integrated omics analyses in identifying shared molecular mechanisms and potential biomarkers.

In summary, our findings show that age, APOE genotype, diet, and sex collectively modulate odor-driven behaviors and memory in mice, with distinct concentrations revealing specific nuances in olfactory preference and exploratory drive. APOE2 mice appeared relatively resilient under high-fat diet conditions, whereas APOE4 carriers displayed patterns indicative higher vulnerability. Memory performance over short-term, long-term, and remote intervals further underscored the interplay between genotype and age, suggesting differential effects on consolidation and retrieval. Elastic MCCA analyses identified key brain connections (notably involving the ventral orbital cortex and primary somatosensory cortex) that underpin odor recognition and associative memory, offering insight into how facilitative and compensatory circuits balance sensory and cognitive demands. Top correlated gene sets between olfactory region and brain were enriched for apoptosis and homeostasis, while in blood, correlated PCs unexpectedly revealed synaptic signaling and oxidative stress pathways—highlighting the potential for blood transcriptomics to reflect key neural processes relevant to Alzheimer’s disease risk. These observations point toward targeted interventions—such as genotype-appropriate dietary strategies or therapies—to preserve or enhance olfactory-based behaviors and overall cognitive health across the lifespan.

Our integrative analysis demonstrates that the molecular basis of olfactory–brain and brain–blood transcriptomic coupling is rooted in shared pathways of apoptosis, synaptic regulation, and metabolic homeostasis. The convergence of these processes across compartments and their association with high-loading genes reinforces their relevance to neural health and disease. These results not only provide mechanistic insight into brain–blood communication but also highlight candidate targets for biomarker discovery and intervention in neurodegenerative disorders.

## Conclusion

Our study demonstrates that olfactory-guided behaviors are sensitive to the combined effects of genetic and environmental risk factors for Alzheimer’s disease. Through integration of behavioral assays, neuroimaging, and transcriptomic profiling, we reveal that early alterations in sensory and cognitive processing are mirrored by distinct changes in brain connectivity and peripheral gene expression. Notably, our results highlight the potential of blood-derived transcriptomic modules—enriched for synaptic, metabolic, and immune-related pathways—to serve as accessible biomarkers reflecting central nervous system vulnerability. The convergence of molecular and systems-level signatures across olfactory and memory circuits underscores the value of integrative approaches in understanding the early mechanisms of neurodegenerative disease and may inform the development of translatable early biomarkers and targeted interventions for AD risk.

